# Single-shot off-axis full-field optical coherence tomography

**DOI:** 10.1101/2022.06.02.494183

**Authors:** Emmanuel Martins Seromenho, Agathe Marmin, Sybille Facca, Nadia Bahlouli, Stephane Perrin, Amir Nahas

**Affiliations:** ICube Research Institute, CNRS - University of Strasbourg, 67085 Strasbourg, France; Department of Hand Surgery, University Hospital of Strasbourg, 67403 Illkirch, France

**Author notes:** (Electronic mail).

## Abstract

Full field optical coherence tomography (FF-OCT) enables high-resolution in-depth imaging within turbid media. In this work, we present a simple approach which combines FF-OCT with off-axis interferometry for the reconstruction of the *en-face* images. With low spatial and temporal coherence illumination, this new method is able to extract an FF-OCT image from only one interference acquisition. This method is described and the proof-of-concept is demonstrated through the observation of scattering samples such as organic and *ex-vivo* biomedical samples.

## I. INTRODUCTION

Full-field optical coherence tomography (FF-OCT) is a noninvasive and label-free imaging technique allowing microscopy imaging in scattering samples^1^. Based on low-coherence interferometry, it selects only the signal from the photons back scattered from the coherence plane (*i.e*., ballistic photons which interfere around the zero optical path difference)^2^. Furthermore, a stack of *en-face* images can then be retrieved, making it possible to reconstruct a three-dimensional amplitude image of biological tissues. An alternative to the full-field direct measurement of the echo time delay of light consists in performing the measurements in the frequency domain by encoding the signal in wavelength^3,4^. However, the acquisition time of the cameras must be adapted, leading to a low signal collection. Furthermore, with a low-spatial-coherence illumination, time-domain FF-OCT has the advantage of reducing the generation of optical aberrations^5,6^ and of eliminating cross talks (*i.e*., interferences between adjacent pixels)^7^. With a typical micrometer resolution^8^, a low speckle dependence^9^ and a high sensibility^10^, FF-OCT has become attractive in many biomedical applications such as optical biopsy in the human gut^11^, the study of intracellular motility in the retina^12^ or high-resolution elasticity measurement in tissues^13,14^.

To reconstruct the two-dimensional *en-face* images at a given axial position within the sample, FF-OCT needs to extract the amplitude from the interference signal. For this purpose, a phase-shifting algorithm is usually implemented using at least three interference acquisitions^15–17^. However, these time-encoded acquisitions do not appear to be well adapted for real-time observation of *in-vivo* tissues. FF-OCT is thus currently limited in its possibility of application to *in-vivo* imaging and clinical transfer.

To address this drawback, methods for single-shot FF-OCT have thus been developed. To our knowledge, there is currently two types of methods described in the literature. The first type is based on light polarization, allowing for the simultaneous recording of the phase-shifted interference patterns^18,19^. Žurauskas *et al*. have proposed a very promising adaption of these polarization-based approaches by illuminating the sample using unpolarized light^20^. The second type is based on off-axis digital holography using low temporally coherent sources^21^. Unlike polarization-based methods, this type of methods requires a spatially coherent illumination source which is not adapted for imaging scattering media because of the cross talk^7^. For this reason, they are usually limited to semi-transparent tissues like the cornea or retina. Swept source off-axis full field OCT approaches were proposed to address this problem by adding membranes^22^ or multimode fibers^23^ to reduce the spatial coherence of the sources. However, the source sweeping requires each image to be temporally coherent. One way to tackle the problem of low spatially coherent illumination in off axis digital holography is to use a diffraction grating as demonstrated in transmission by Slabý *et al*. for quantitative phase imaging ^24^.

In this study, we demonstrate a new approach in FF-OCT that enables single-shot acquisitions through the concept of off-axis digital holography. To our knowledge, off-axis interferometry with a low spatially and temporally coherence illumination is for the first time employed in order to perform tomography in scattering samples. To record the full-field holograms with a low coherence light source, a diffraction grating and a series of lenses are introduced in the illumination part, making it possible both to tilt and to rotate the coherence plane, respectively. This method has the advantage of being robust (*i.e*., motion artefacts are avoided), simple and fast (*i.e*.,real time observation can be performed). Indeed, by being only limited by the frame rate of the camera and the computational speed, the reconstruction time of an *en-face* image from a single-shot acquisition is considerably improved.

## II. METHOD

The system consists of a FF-OCT system where the illumination part has been enhanced to perform full-field off-axis interference measurements. Indeed, as illustrated in Fig. 1(a), the illumination part provides two spatially-separated beams instead of the usual single one. For this purpose, the collimated incident beam from a LED (Thorlabs M660L4, λ_0_ = 660 nm, FWHM = 20 nm) is diffracted by a grating (Thorlabs GT25-03, Λ = 300 l/mm) in order to generate the two illumination beams. The period of the grating grooves matches with the shift in the frequency domain. Moreover, the coherence plane of the first-diffraction-order beam is then tilted by an angle *α* relative to the propagation direction^25^, while the zero-order beam is not affected. According to the dispersion relation, the angle *α* is equal to the diffraction angle. Then, an achromatic lens collects the two beams and forms two images of the light source in its focal plane. Afterwards, each beam passes separately through an afocal optical system (achromat doublet lenses, f’ = 75 mm), in order to flip the coherence planes individually^26^ before being transmitted and reflected by a beam splitter cube (ratio of 50:50). Without flipping the coherence plane to compensate the double pass through the microscope objectives, the interference pattern would be narrowed by the coherence length. The theoretical principle has previously been described for digital holography in transmission mode^24^. The single-shot FF-OCT system is mainly based on this innovative combination of a diffraction grating with a couple of lenses, allowing for a round trip of the beam in reflection mode.

**FIG. 1.**
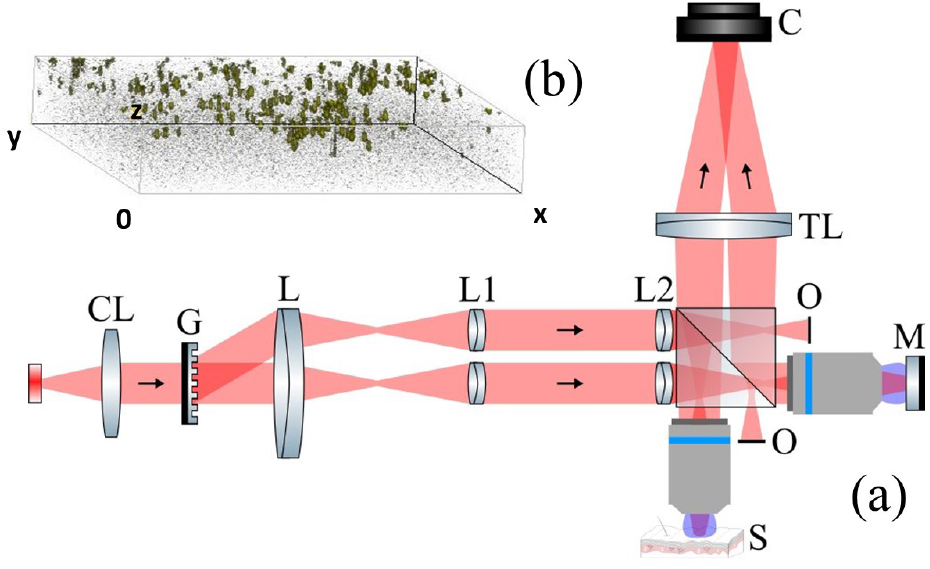
(a) Layout of the off-axis OCT system for retrieving *en-face* amplitude full-field distributions in a single shot. CL, collimating lens. G, grating. L, lens. L1 and L2, afocal system. O, obstruction. M, mirror. MO, microscope objective. S, sample. TL, tube lens. C, camera. (b) Volume reconstruction of gold nanoparticles immersed in agarose gel on which a cover slice is deposited. The field of view is cropped to be more readable (200 *μm* × 200 *μm*). Stack of 80 off-axis interferograms with Δ*z* =1 *μ*m.

The transmitted zero-order beam becomes the reference beam and the reflected first-order beam, the object beam (the two other beams are dumped). Based on a Linnik configuration, two similar water-immersion microscope objectives (Olympus, ×10, NA = 0.3) are introduced in the reference and the object arms at the back focal plane of the last lens L2. Finally, the back-reflected beams from the reference mirror and the sample are combined and directed on a camera (Adimec Q-2HFW, 1440 × 1440 pixels, pitch = 12 *μ*m, 500 frames/s) using a tube lens (*f′* = 500 mm). The angle between the two beams is again *α*, thus making it possible to record a full-field off-axis hologram at a given depth^27^. The intensity distribution of the recorded interferogram can be expressed as:

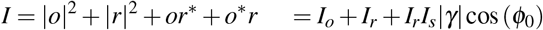

where *o* and *r* are the complex electric fields of the object and the reference arms, respectively. * represents the complex conjugate. The relation consists of the complex field of the sample (*i.e*., the amplitude |*γ*| term and the phase *ϕ*_0_ term). The transverse amplitude distribution which depends on the sample reflectivity, is extracted using an angular spectrum algorithm^28^. In order to quantify the performance of the FF-OCT system, 500-nm-diameter gold nanoparticles (Sigma Aldrich®) were immersed in agarose gel and imaged with the system. A part of the reconstructed volume is presented in Fig. 1(b). A thin cover glass flattens the sample, yielding an aggregate of nanoparticles at the top of the sample. The 3D point spread function is estimated using an isolated nanoparticle, resulting in a lateral resolution of about 1.7 μm and an axial resolution of about 6.9 *μ*m. The lateral field of view reaches 620 *μm* and the lateral magnification, around ×28. To acquire a volumetric image of the sample, a stack of temporally-encoded off-axis interferograms was recorded by axial translation of the sample. As in classical FF-OCT, the imaging depth is limited by scattering in biological tissues and optical aberrations, and reaches about 100 —200 *μm* in the visible range. Assuming shot-noise-limited measurements, the sensibility is estimated at about 75 dB (corresponding to a minimal reflectivity of 3.4 × 10^−8^).

## III. RESULTS

### A. *En-face* amplitude images of scattering samples

To validate the proof-of-concept, amplitude images of *ex-vivo* scattering samples have been acquired (*cf*. Fig. 2). All *en-face* amplitude distributions are reconstructed using only one interferogram acquisition. The amplitude images are obtained on plants and *ex-vivo* animal tissues. Fig. 2(a,b) represent *en-face* amplitude images from samples of spider-plant leaf (*chlorophytum comosum*) and ficus leaf (*ficus benjamina*) respectively. It is therefore possible to obverse the particular geometry of the spider-plant leaf cells^29^ (*cf*. Fig 2(a)) and the smaller ficus leaf cells, where the black round structure corresponding to lithocysts is shown (*cf*. Fig 2(b)). Freshly excised tissues were also observed through the single-shot FF-OCT system such as mouse brain (*cf*. Fig. 2(c)) and chicken skin (*cf*. Fig. 2 (d)). The myelins fibers in the mouse brain and collagen fibers of the chicken skin are clearly visible.

**FIG. 2.**
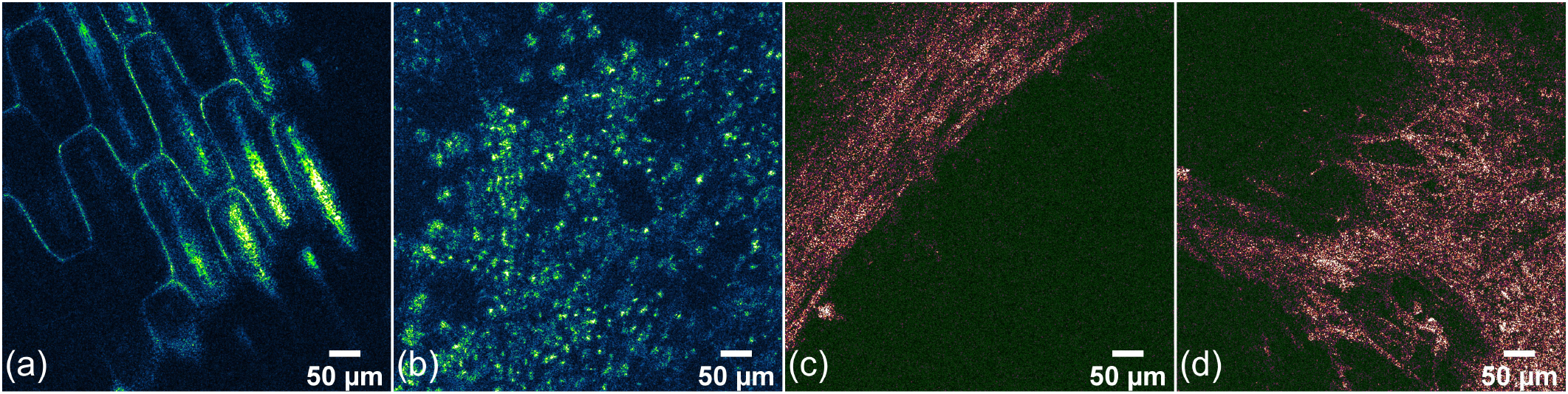
*En-face* amplitude distribution images (in false colors) of spider-plant leaf (a), ficus leaf (b), mouse brain (c), and chicken skin at a depth of 22 *μm*, 38 *μm*, 30 *μm*, and 44 *μm* respectively.

### B. Images comparison between commercial FF-OCT and single-shot FF-OCT

A comparison between single-shot FF-OCT and a commercial FF-OCT (Acquyre Biosciences©) was carried out to validate the single-shot FF-OCT approach proposed in this manuscript. Fig. 3 represents *en-face* amplitude images of a spider-plant leaf sample. Fig. 3(a) was obtained with the commercial FF-OCT (Acquyre Biosciences©) and Fig. 3(b) with the single-shot FF-OCT approach. Both images were acquired in similar conditions (i.e., 40 frames average and close to camera saturation). Qualitatively, both images are similar with typical signal to noise ratio (SNR) (measure on cell walls) of 5.4 for the commercial FF-OCT and 18.9 for the single-shot FF-OCT. The higher SNR for the single-shot FF-OCT could be explained by the larger coherence length of the source compared to the commercial system.

**FIG. 3.**
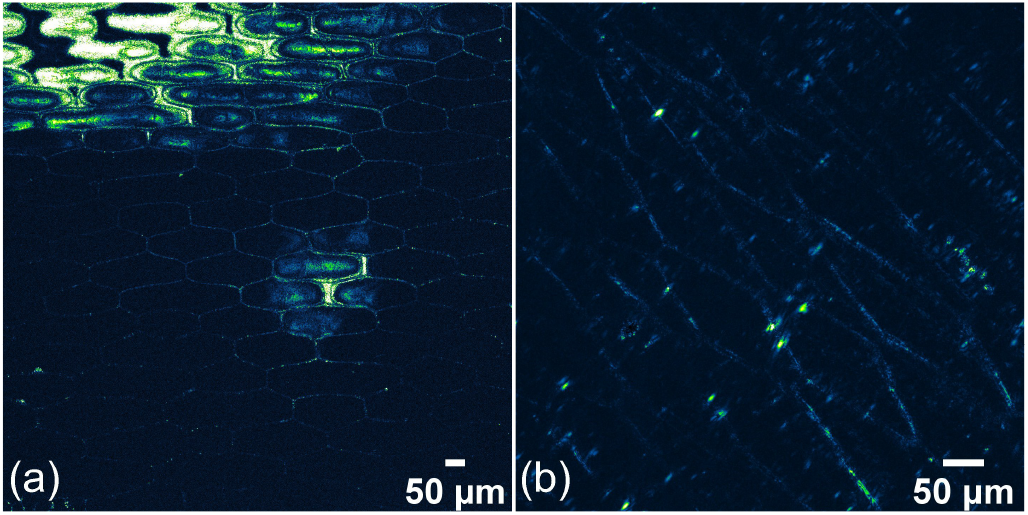
*En-face* amplitude images (in false colors) of spider-plant leaf at a depth of 55 μm. (a) Image from commercial FF-OCT, the field of view is 1266×1266 μm^2^. (b) Image from single-shot FF-OCT, the field of view is 620 × 620 *μm*^2^.

In both Fig. 3(a) and Fig. 3(b) the typical geometry of spider plant epidermal cells is recognizable. The major difference between the two images is the field of view, which has to be limited in the off-axis configuration to keep diffraction limited performances.

## IV. DISCUSSION

In comparison with classical FF-OCT, the lateral field of view appears to be reduced by a factor of two due to the fringe sampling. Moreover, this FF-OCT system leads to around a quarter of the usual light budget. For this, dark-field configuration could enhance the intensity collection^30^. In addition, this method is not only able to extract the *en-face* amplitude from one interferogram, but also to provide a direct access to the phase distribution, thus giving an access to new contrast modalities such as shearwave elastography^31^, photo-thermal contrast and Doppler imaging properties of living samples.

## V. CONCLUSION

In conclusion, the principle of single-shot FF-OCT using a low spatially and temporally coherence illumination is presented for the first time. The method is validated by imaging in-depth *ex-vivo* biological samples. This approach consists of a relatively simple experimental arrangement where only one interferogram is collected. A digital-holography-based algorithm then makes it possible to retrieve the *en-face* amplitude image at a given depth from the single acquisition. This method provides a significant enhancement in FF-OCT for *in-vivo* imaging where, usually, the sample motions strongly prevent phase-shifted acquisitions without limiting the dynamic.

## ACKNOWLEDGMENTS

The authors thank Aurore de Cauwer and Chrystelle Po for providing the biological samples, Jean Marie Chassot for providing the FF-OCT commercial (Acquyre Biosciences©) images, Amandine Elchinger and Vincent Maioli for their constructive inputs in writing the manuscript, and Jesse Schiffler for his help for the experimental setup.

## DATA AVAILABILITY

Data underlying the results presented in this paper are not publicly available at this time but may be obtained from the authors upon reasonable request.

## References

1 A. Dubois and A. Boccara, “Full-field optical coherence tomography,” in Optical coherence tomography, edited by W. Drexler and J. Fujimoto (Springer, 2008).

2 R. A. Leitgeb, “En face optical coherence tomography: a technology review,” Biomedical Optics Express 10, 2177–2201 (2019).

3 C. Gorecki, J. Lullin, S. Perrin, S. Bargiel, J. Albero, O. Gaiffe, J. Rutkowski, J. M. Cote, J. Krauter, W. Osten, W.-S. Wang, M. Weimer, and J. Froemel, “Micromachined phase-shifted array-type mirau interferometer for swept-source oct imaging: design, microfabrication and experimental validation,” Biomedical Optics Express 10, 1111–1125 (2019).

4 I. F. Almog, F.-D. Chen, S. Senova, A. Fomenko, E. Gondard, W. D. Sacher, A. M. Lozano, and J. K. S. Poon, “Full-field swept-source optical coherence tomography and neural tissue classification for deep brain imaging,” Journal of Biophotonics 13, e201960083 (2020).

5 P. Xiao, V. Mazlin, K. Grieve, J.-A. Sahel, M. Fink, and A. C. Boccara, “In vivo high-resolution human retinal imaging with wavefront-correctionless full-field oct,” Optica 5, 409–412 (2018).

6 V. Barolle, J. Scholler, P. Mecê, J.-M. Chassot, K. Groux, M. Fink, A. C. Boccara, and A. Aubry, “Manifestation of aberrations in full-field optical coherence tomography,” Optics Express 29, 22044–22065 (2021).

7 B. Karamata, P. Lambelet, M. Laubscher, R. P. Salathé, and T. Lasser, “Spatially incoherent illumination as a mechanism for cross-talk suppression in wide-field optical coherence tomography,” Optics Letters 29, 736–738 (2004).

8 A. Dubois, L. Vabre, A.-C. Boccara, and E. Beaurepaire, “High-resolution full-field optical coherence tomography with a linnik microscope,” Applied Optics 41, 805–812 (2002).

9 S. M. S. Kazmi, R. K. Wu, and A. K. Dunn, “Evaluating multi-exposure speckle imaging estimates of absolute autocorrelation times,” Optics Letters 40, 3643–3646 (2015).

10 A. Dubois, K. Grieve, G. Moneron, R. Lecaque, L. Vabre, and C. Boccara, “Ultrahigh-resolution full-field optical coherence tomography,” Applied Optics 43, 2874–2883 (2004).

11 L. Quénéhervé, R. Olivier, M. J. Gora, C. Bossard, J.-F. Mosnier, E. Benoit a la Guillaume, C. Boccara, C. Brochard, M. Neunlist, and E. Coron, “Fullfield optical coherence tomography: novel imaging technique for extemporaneous high-resolution analysis of mucosal architecture in human gut biopsies,” Gut 70, 6–8 (2021).

12 J. Scholler, K. Groux, O. Goureau, J.-A. Sahel, M. Fink, S. Reichman, C. Boccara, and K. Grieve, “Dynamic full-field optical coherence tomography: 3d live-imaging of retinal organoids,” Light: Science & Applications 9, 140 (2020).

13 A. Nahas, M. Tanter, T.-M. Nguyen, J.-M. Chassot, M. Fink, and A. C. Boccara, “From supersonic shear wave imaging to full-field optical coherence shear wave elastography,” Journal of Biomedical Optics 18, 1 – 9 (2013).

14 O. Thouvenin, C. Apelian, A. Nahas, M. Fink, and C. Boccara, “Full-field optical coherence tomography as a diagnosis tool: Recent progress with multimodal imaging,” Applied Sciences 7 (2017), 10.3390/app7030236.

15 E. Beaurepaire, A. C. Boccara, M. Lebec, L. Blanchot, and H. Saint-Jalmes, “Full-field optical coherence microscopy,” Optics Letters 23, 244–246 (1998).

16 W. Y. Oh, B. E. Bouma, N. Iftimia, S. H. Yun, R. Yelin, and G. J. Tearney, “Ultrahigh-resolution full-field optical coherence microscopy using ingaas camera,” Optics Express 14, 726–735 (2006).

17 J. Na, W. J. Choi, E. S. Choi, S. Y. Ryu, and B. H. Lee, “Image restoration method based on hilbert transform for full-field optical coherence tomography,” Applied Optics 47, 459–466 (2008).

18 C. Dunsby, Y. Gu, and P. M. W. French, “Single-shot phase-stepped wide-field coherence-gated imaging,” Optics Express 11, 105–115 (2003).

19 K. Creath and G. Goldstein, “Dynamic quantitative phase imaging for biological objects using a pixelated phase mask,” Biomedical Optics Express 3, 2866–2880 (2012).

20 M. Žurauskas, R. R. Iyer, and S. A. Boppart, “Simultaneous 4-phase-shifted full-field optical coherence microscopy,” Biomedical Optics Express 12, 981–992 (2021).

21 H. Sudkamp, P. Koch, H. Spahr, D. Hillmann, G. Franke, M. Münst, F. Reinholz, R. Birngruber, and G. Hüttmann, “In-vivo retinal imaging with off-axis full-field time-domain optical coherence tomography,” Optics Letters 41, 4987–4990 (2016).

22 P. Stremplewski, E. Auksorius, P. Wnuk, Ł. Kozon, P. Garstecki, and M. Wojtkowski, “In vivo volumetric imaging by crosstalk-free full-field oct,” Optica 6, 608–617 (2019).

23 E. Auksorius, D. Borycki, P. Wegrzyn, I. Žičkienė, K. Adomavicius, B. Sikorski, and M. Wojtkowski, “Multimode fiber as a tool to reduce cross talk in fourier-domain full-field optical coherence tomography,” Optics Letters 47, 838–841 (2022).

24 T. Slabý, P. Kolman, Z. Dostál, M. Antoš, M. Lošt’ák, and R. Chmelík, “Off-axis setup taking full advantage of incoherent illumination in coherence-controlled holographic microscope,” Optics Express 21, 14747–14762 (2013).

25 S. Akturk, X. Gu, E. Zeek, and R. Trebino, “Pulse-front tilt caused by spatial and temporal chirp,” Optics Express 12, 4399–4410 (2004).

26 P. Bouchal, R. Chmelík, and Z. Bouchal, “Phase of white light and its compatibility to the optical path,” Optics Express 29, 12398–12412 (2021).

27 T. Balcunas, A. Melninkaitis, A. Vanagas, and V. Sirutkaitis, “Tilted-pulse time-resolved off-axis digital holography,” Optics Letters 34, 2715–2717 (2009).

28 E. Cuche, F. Bevilacqua, and C. Depeursinge, “Digital holography for quantitative phase-contrast imaging,” Optics Letters 24, 291–293 (1999).

29 W. Charlton, “Differentiation in leaf epidermis of chlorophytum comosum baker,” Annals of Botany 66, 567–578 (1990).

30 E. Auksorius and A. C. Boccara, “High-throughput dark-field full-field optical coherence tomography,” Optics Letters 45, 455–458 (2020).

31 A. Marmin, S. Catheline, and A. Nahas, “Full-field passive elastography using digital holography,” Optics Letters 45, 2965–2968 (2020).

